# Model based normalization improves differential expression calling in low-depth RNA-seq

**DOI:** 10.1101/013854

**Authors:** Pavel N. Zakharov, Alexey A. Sergushichev, Alexander V. Predeus, Maxim N. Artyomov

## Abstract

RNA-seq is a powerful tool for gene expression profiling and differential expression analysis. Its power depends on sequencing depth which limits its high-throughput potential, with 10-15 million reads considered as optimal balance between quality of differential expression calling and cost per sample. We observed, however, that some statistical features of the data, e.g. gene count distribution, are preserved well below 10-15M reads, and found that they improve differential expression analysis at low sequencing depths when distribution statistics is estimated by pooling individual samples to a combined higher-depth library. Using a novel gene-by-gene scaling technique, based on the fact that gene counts obey Pareto-like distribution^1^, we re-normalize samples towards bigger sequencing depth and show that this leads to significant improvement in differential expression calling, with only a marginal increase in false positive calls. This makes differential expression calling from 3-4M reads comparable to 10-15M reads, improving high-throughput of RNA-sequencing 3-4 fold.

## Introduction

RNA-Seq is an actively used technique for transcriptome profiling, providing a powerful alternative to microarrays due to several strong advantages over it. First, RNA-Seq does not require existing information about the genome sequence, and thus allows analysis of transcription, measuring unknown trnascripts including novel non-coding genome regions. Second, RNA-Seq introduces less background noise, avoiding the microarray’s probe cross-hybridization noise. Finally, since there is no upper limit for quantification in RNA-Seq, it gives a much wider dynamic range of detection in comparison with the microarray method^2^, which has both saturation and background limitations. All these benefits of RNA-Seq make it very attractive for the key task in functional genomics – identifying differential expression (DE).

However, the ability to find differential expression is strongly influenced by the sequencing depth – the deeper sequencing, the higher the statistical power to detect differentially expressed genes^3^. On the other hand, to achieve maximum power within the same budget, one should find a trade-off between sequencing depth and number of samples sequenced. Although the power to detect differentially expressed genes increases with both the increased number of biological replicates and increased sequencing depth, sequencing deeper a certain number of reads in each sample generates the diminishing returns for the power of detecting DE genes. According to Liu et al., 2013^4^, the increase in number of DE genes with sequencing depth gives lower outcome after 10-15 million (10-15M) reads. Furthermore, for practical reasons one usually does not go beyond duplicates or triplicates in the experimental design while a substantial increase in power with increase of replication occurs independently of sequencing depth up to 5-7 samples^4^.

Given the current instrumental throughput capabilities, such sequencing depth still makes RNA-seq experiments prohibitively expensive for high-throughput studies. One avenue to increase throughput of RNA-seq based studies is to compensate for decreases in sequencing depth by leveraging additional information present in RNA-seq data. In this work, we show that global read coverage distribution provides the source of such information that can be leveraged to improve detection of differential expression at the sequencing depths of 3-5 million reads to match that of 10-12 million reads sequencing depth with only minor loss of specificity.

## Results

Consistent with previous estimates^4^, we have also observed that 10-15 million reads is the optimal sequencing depth using the common benchmark RNA-seq dataset from Bottomly et al^5^, comparing downsampled datasets from two different mouse strains, B6 and D2, originally sequenced at ∼30M reads per sample. As one can see from Fig. 1a, most of the differentially expressed genes are called in the libraries of sequencing depth of 10-15 M, and with further depth increase, number of detected DE genes increses only decrementally (using genes with adjusted p-value below 0.05 at 30M depth in 3 replicates vs 3 replicates comparison as a set of truly DE genes, further referred as the **verification set** here). Furthermore, the number of false negative(FN) DE genes decreases almost twofold upon increasing depth from 1M to 10M, while the p-values of these FN genes drastically increase. This means that only less significant DE genes are lost when increasing sequencing depth beyond ∼10M reads. Overall, this confirms that sequencing depth of ∼10-15M represents the current “golden ratio” point for RNA-seq in terms of balancing multiplexing and quality of differential expression calling.

**Figure 1.**
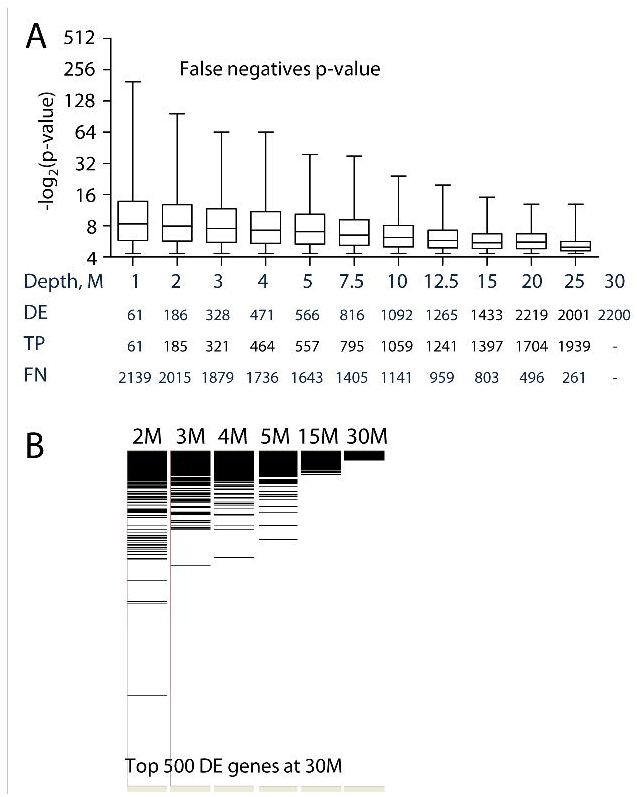
Most of the significant differentially expressed genes are called in the libraries of sequencing depth of 10M-15M. Result of 3 vs 3 samples comparisons from the Bottomly et al. dataset.

A. ‒Log2(adj. p-value) of FN genes. P-values obtained from 30M sequence depth in 3 vs 3 sample comparison. Statistical power of not detected DE genes decrease mostly to 10M reads per sample and then after 10M-12.5M doesn’t change much.
B. 500 most significantly expressed genes from the verification set (defined above) mostly preserve their ranking at sequencing depths below 10M. Black lines correspond to top 500 genes with lowest p-values from the verification set. All genes are ranked top down by adjusted p-value increase. Enrichment bars obtained using GSEA software (http://www.broadinstitute.org/gsea). DE genes have mostly the highest statistical power and are presented at the top of the ranked lists. But it is not clear how to call directly DE genes since statistical power of differential expression goes down with decreasing number of reads.

While ability to determine differential expression declines rapidly, not all the information is lost at an equal pace when reducing sequencing depth below 10M. In fact, as seen in Fig. 1b, relative ranking of DE genes is mostly preserved at as low a sequencing depth as 2M per library, even though accurate calling of differential expression is not directly feasible at 2M, due to the loss of statistical power (see, for example, table in the panel Fig. 1a). Thus, we have asked if it was possible to leverage the information content that is preserved below 10M depth to recover information about DE genes at the lower sequencing depth.

To address this question we have examined the statistical properties of RNA-seq datasets to find the most conserved characteristics, starting from gene level distribution of sequencing reads. It has been shown recently that the double Pareto lognormal (dPLN) fitting provides a very accurate approximation of the gene expression reads distribution for RNA sequencing data^1,6^. Briefly, its main improvement over the lognormal fitting is that it better describes the end segments of the distribution corresponding to both low and high numbers of reads. Since for the random value X distributed with the dPLN law, log(X) follows the Normal Laplace distribution (NL), we used the natural logarithm of the reads and the NL density function for fitting^1,7,8^. Indeed, we observed that the NL density function accurately fits gene level distribution of sequencing reads at both low and high sequencing depths (Fig 2a-b, Supplementary Fig. 1a-b).

**Figure 2.**
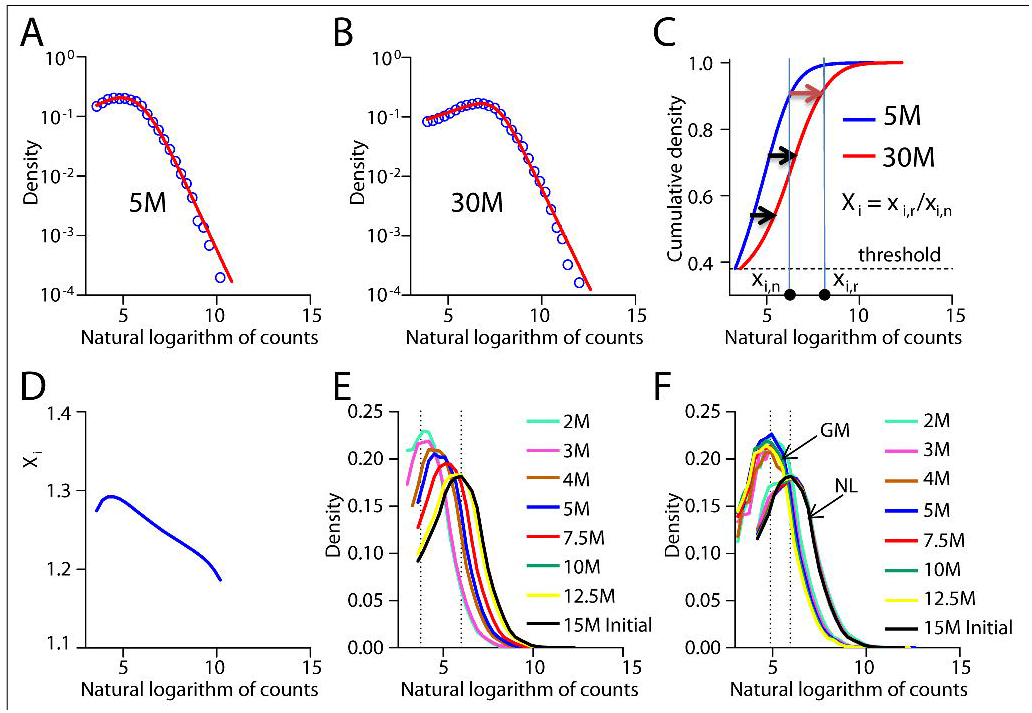
Fitting and transformation procedure of RNA-Seq distributions. We consider here as an example the two B6 mouse strain sequencing data with 30M and downsampled from it 5M sequencing depth

A. Distributions for the RNA-Seq datasets with 30M(right) and downsampled 5M(left) sequencing depth. Logarithmed distributions were fitted with Normal-Laplace functions after cut off was applied. Y-axes are in log scale.
B. Transformation procedure. Two integral distribution curves correspond the two differential distributions at A. Threshold was applied by cutting off the curves below the certain y-value (dashed line). The 5M is being transformed (normalized) to 30M. Coefficients of normalization Xi are individual for each gene.
C. Dependence of distribution normalization coefficients from C. on the gene expression levels (natural logarithm of reads per gene). Coefficients are nonuniform and change across expression levels.
D. Initial not transformed RNA-Seq reads distributions for 2M-15M sequence depth. Going from the 2M to 15M the more distinct peak is observed.
E. Geometric mean transformed distributions from D.
F. Normal-Laplace transformed distributions from D. The peak of the normalized distributions is at about 5.8 as opposed to 4.9 at F., which is a result of normalization to the largest library distribution instead of averaging all distribution datasets. Each curve at D-F corresponds to average of 3 downsampling repeats from one biological replicate. The y-axis corresponds to the normalized density values and the x-axis to the natural logarithm of expression calls number.

In the context of microarray studies, typical shape of signal distribution is used to normalize the data; while in the context of RNA-sequencing data, normalization has played little role other than equilibrating sequencing depth between different samples. Most of differential expression methods are traditionally more concerned with accurate statistical representation of noise impact, rather than the shape of read distribution. Typical sample normalization includes only simple rescaling to average library depth, using a common normalization factor for all genes in individual libraries, such as geometric mean (GM) normalization employed in DESeq^9^, DEXSeq^10^ and DESeq2^11^. We asked if a model-based approach with explicit descriptions of gene counts distribution (e.g. based on double Pareto lognormal) could improve accuracy of differential expression calling in the regimes of low sequencing depths. A major potential advantage of using explicit distributions is the ability to model RNA-Seq dataset at higher sequencing depths, effectively re-scaling the data from 2-5M towards 10-15M in a statistically and biologically meaningful way.

To investigate this possibility in details, we devised the following normalization procedure, illustrated by scaling 5 million read depth dataset (obtained through downsampling) towards its original 30 million reads deep sequencing of B6 mouse cells from Bottomly et al data^5^ (Fig.2a-b). First, a cumulative distribution function is built for both datasets and a cutoff is introduced to the 5M dataset to avoid considering non-expressed genes with zero or few reads. Considering cumulative distribution curves, this cutoff value is re-scaled for deeper sequencing distribution by finding a point with the same y-axis value, reflecting the fact that area under the curve (i.e. cumulative probability of non-expressed genes) should be preserved between two distributions (Supplementary Fig. 2). Next, we estimate the parameters of corresponding Pareto distributions for each dataset by plotting a histogram of gene counts (Fig. 2a-b,e), bins are depicted with empty circles. Given explicit functions for each distribution, we scale 5M dataset on a gene-by-gene basis. The normalization approach consists of multiplying each count from the initial distribution by the coefficient X_i_ = x_i,r_/x_i,n_, where x_i,n_ is the x-coordinate corresponding to the i^th^ gene read count in the non-normalized distribution and x_ir_ is x-coordinate at the 30M reference curve with the same y-coordinate as x_ir_ (Fig. 2c). Notably, coefficients X_i_ are nonequal across the expression levels in our normalization approach (Fig. 2d).

The main difference between our NL normalization approach and the geometric mean (GM) transformation is that the first one normalizes distributions to the reference with the highest depth of sequencing, while the GM approach comes up with the averaged distribution (see dashed lines on Fig.2 e-f. showing the peak positions.). Hence the resulting distributions will correspond to the deepest one, in the case of NL, and to the mean one, in the GM case (Fig. 2e-f). In other words, the NL transformation takes advantage of the samples with the highest sequencing depth and thus has potential to increase DE detection power as we show below. Importantly, the GM normalization uses single normalization factor per one distribution leading to distortion at the ditribution tails, while NL method leverages explicit model effectively allowing for number of normalization factors equal to the number of genes in the dataset. As a consequence, the NL transformation gives more tight fits than the GM normalization (Fig. 2f).

To evaluate the effect of the NL transformation towards higher depth on the quality of the differential expression calling, we have performed differential expression analysis for 3 B6 vs 3 D2 replicates per sample, for data by Bottomly et al.^5^ for both initial and the NL transformed datasets at different levels of downsampling. Data downsampled to different depths from 2 to 10 million reads were NL renormalized to known 15M dataset distribution by the procedure described above. The DE genes were found via DESeq2^11^ for each downsampled sequencing depth with adjusted p-value threshold of 0.05. We then calculated sensitivity, precision, specificity, and accuracy of differential expression calls using verification set defined above (see also Methods).

As expected, the number of detected DE genes increased with the growth of sequencing depth for both NL transformed and GM normalized data (Fig. 3a). Importantly, the number of differentially expressed genes called upon NL transformation was several fold higher in the cases of low depth libraries, hence dramatically improving the amount of differnetially expressed genes. To see if the increase was due to inclusion of true positive DE genes previously lost due to insufficient statistical power, we computed sensitivity and precision at each level of downsampling. Indeed, NL transformed data showed higher sensitivity in comparison with not normalized (Fig. 3b): for NL transformed 2M data we got roughly the same amount of DE genes as for 7.5M not transformed dataset and for 5M NL transformed data result was similar to between 12.5M - 15M not transformed dataset. Such increase in sensitivity (2.4 times for 2M and 1.8 times for 5M) was accompanied with only mild drop of precision – about 10-20%. Accuracy and specificity metrics were almost unchanged (Supplementary Fig. 3a-b). Thus, we conclude that provided information about gene count distribution at higher sequencing depths, one can drastically increase sensitivity of the DE detection while maintaining good precision.

**Figure 3.**
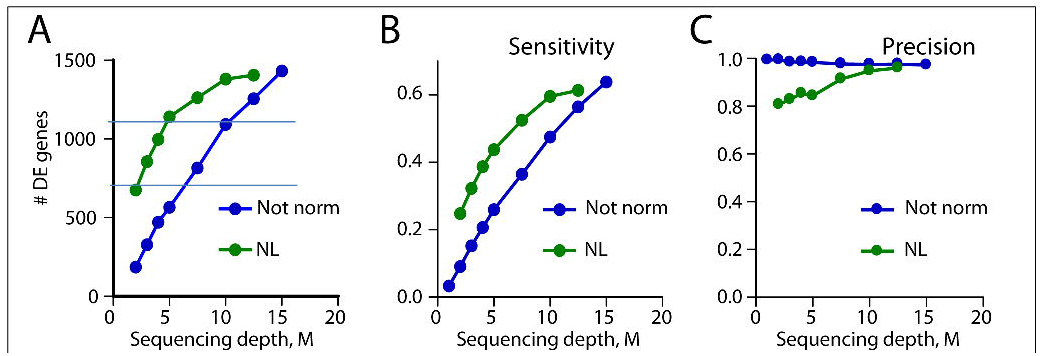
Benchmark of DE calls, sensitivity and precision for data normalized to 15M reference distribution with 3 vs 3 biological replicates comparison. The verification true positive set was the DE genes detected at ∼30M library size for 3 vs 3 biological replicates comparison. The DE genes at the current sequencing depth were defined from comparisons of 3 vs 3 samples from the Bottomly et al.^5^ with adjusted p-values less than 0.05. Green lines are result of NL normalization, green lines are for not normalized datasets.

A. Dependence of DE calls on sequencing depth for NL normalized and not normalized data. Roughly the same number of DE genes as at 10M was obtained with only 5M NL transformed data, normalization of 2M library yields the amount of DE detected genes as at 5M – 7.5M depth with not normalized datasets.
B. – sensitivity and precision (see definition in text). Very similar increase in sensitivity occured as in DE detection on A, while drop in sensitivity from 2M to 5M was about 20%.

However, for the actual high-throughput experiment with 20-30 samples per lane, the reference "high depth distribution" is not known. To reconstruct a reference distribution one can use a “pooling” approach, summing up reads for each gene through replicates and samples. In our case, we summed up reads for 2 samples with 2 replicates per each sample (Fig. 4a). For example, with 3M sequencing depth we gained effectively 12M depth reference distribution under the assumption that the majority of the genes are not differentially expressed. Then, we NL normalized each sample separately to the pooled reference distribution. As the result in 2 vs 2 replicates per sample comparison, the number of DE genes increased along with the sensitivity, while the precision dropped mildly (Fig. 4 b-d, Table 1) and accuracy, specificity did not change much (Supplementary Fig. 5). Hence, with reference distribution reconstructed through sample pooling we see a significant improvement in the number of found TP genes with a marginal loss in precision.

**Figure 4.**
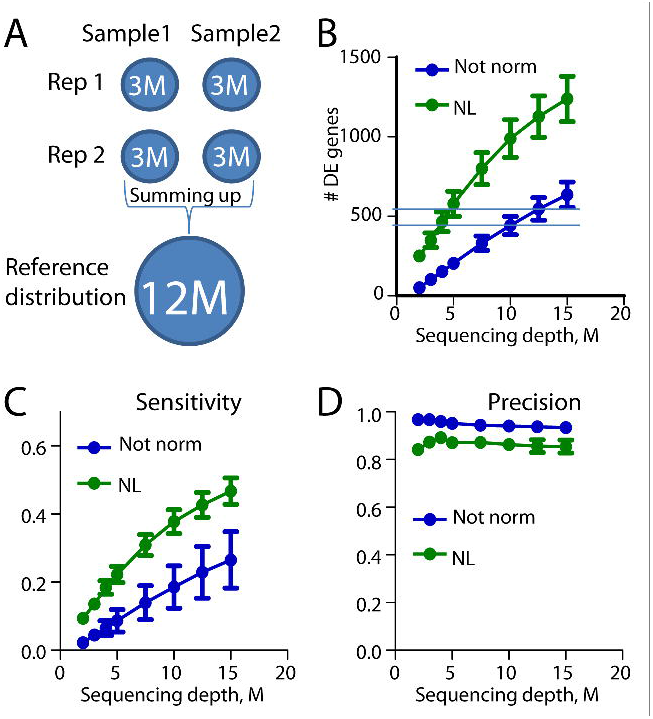
Benchmark of DE calls, sensitivity and precision for data normalized to pooled reference distribution with 2 vs 2 biological replicates comparison. To improve high-throughput, many samples can be sequenced to lower depth and then information from all samples and replicates can be combined together to obtain more information about DE expression at a given sequencing depth. Bars mean +/- SEM. Green lines – NL normalized and blue lines – not normalized data.

A. Schematic representation of the pooling approach. Two biological replicates per each of two samples were summed up so that the counts corresponding to the same genes were combined together. The resulting “pooled” distribution was used as a reference distribution. For example, starting with 3M sequencing depth and 2 replicates per each of two samples the one can obtain effectively 12M reference distribution.
B. Number of detected DE genes for 2 vs 2 biological replicates comparison. The amount of detected at 10M-12.5M DE genes can be achieved with 4M-5M normalized data.
C. and D. – sensitivity and precision. The normalization procedure gave the same result in termes of sensitivity at 2M-5M as the one would get at 5M-10M with initial data.

To pictorially represent dramatic improvement due to NL normalization, we plotted Venn diagrams of intersections between DE gene sets found in normalized and initial data, along with the verification set at low sequencing depths on Fig. 5. The dark brown area shows TP genes represented both in non-normalized and NL transformed data (preserved genes), the bright red region depicts the TP genes found as a result of the transformation procedure in addition to those discovered with non-transformed data. So for the 2 vs 2 sample comparison the transformation gives two times more discovered TP genes at 15M and an almost five times increase in TPs for 2M depth. The number of FP genes found with transformed data (shown in light blue) is much lower than the number of additionally found TP genes. The yellow region depicts the number of lost TP genes after transformation. Additional improvement can be gained by combining DE genes found with and without normalization. It means recovering TP genes lost in the transformation procedure (yellow region), with very low addition of FP genes found with non-transformed data. Detailed information about 2 vs 2 DE analysis with means and errors is given in Supplementary Table 1. To summarize, 4-5M sequencing depth with transformation gives sensitivity very close to 10M for non-transformed data both with 2 vs 2 and 3 vs 3 (see Fig. 4, Supplementary Fig. 4, Supplementary Tables 1-2) replicates comparison, which is usually considered an appropriate sequencing depth for DE analysis.

**Figure 5.**
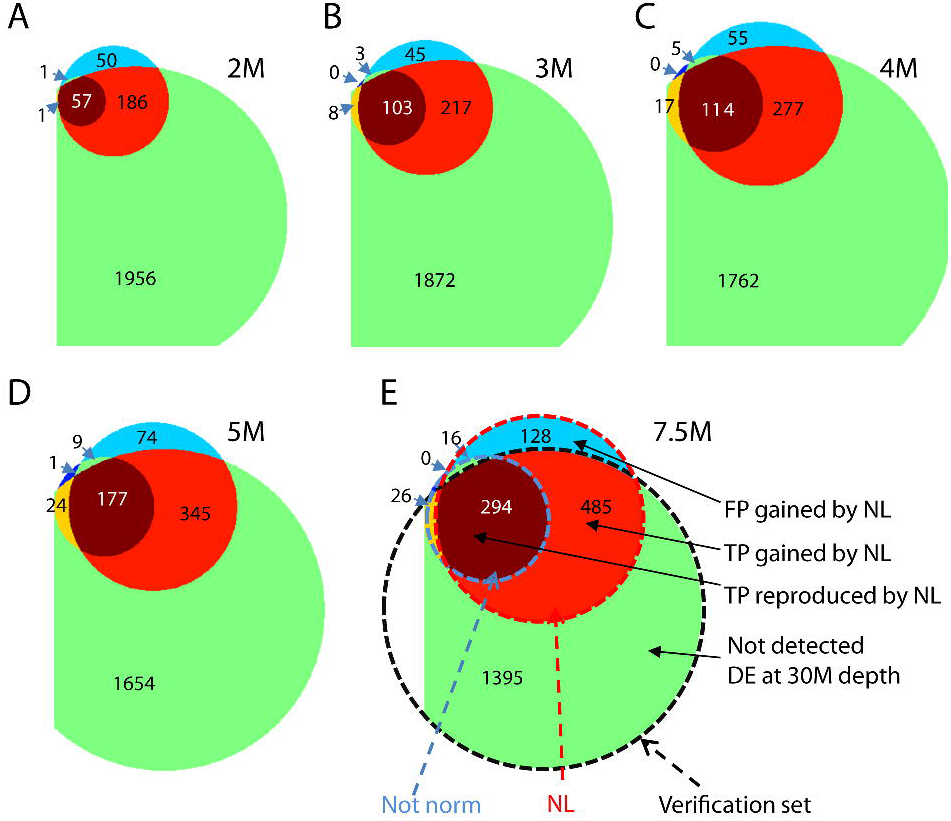
Venn diagrams of intersections between verification gene set, DE genes obtained from 2 vs 2 replicates comparison of not normalized samples and the ones normalized with the pooling approach. Diagrams were plotted for 2M-7.5M sequencing depth. Light blue region represents FPs gained by the NL normalization, dark bright red – TPs gained by NL normalization, dark brown – TPs reproduced by NL normalization, the biggest green region – not detected by both methods DE genes, yellow – DE genes detected without normalization and not found after normalization was applied. After normalization we obtained ∼4 times more TPs at 2M and 2.4 times more at 5M. Almost all TPs found without normalization were reproduced with our approach, while number of gained FPs was comparably low: ∼1/2 at 2M and ∼1/5 at 5M of gained TPs via normalization. See Table 1 in addition to the Venn diagrams.

## Discussion

Currently, the high-thoughput potential of RNA-sequencing relies heavily on both the depth of sequencing and the number of biological replicates per condition. Here we present a computational strategy that leverages information about gene count distribution in order to improve differential expression calling at low sequencing depths. By using additional information about distribution of sequencing gene calls, one can re-scale a RNA-Seq dataset towards higher depth, compared to the one actually achieved in the experiment. As we show, such normalization can increase sensitivity of the sequencing with 4M-5M libraries depth to become comparable to 10M-15M depth with only a mild drop in precision. This approach will be very useful in the high-throughput applications of RNA-sequencing.

## Online Methods

### Libraries alignment, down-sampling and DE genes detection

We used RNA-Seq data from Bottomly et al. obtained for two different mouse strains cells B6 and D2 with 3 biological replicates per sample, SRA026846 (B6: SRR099237, SRR099238, SRR099239 and D2: SRR099240, SRR099241, SRR099242), ∼30M reads in each of 6 libraries. The libraries were single-end reads of 70 bp and were evaluated to be unstranded using CollectRnaSeqMetrics utility from Picard tools. All libraries were aligned to GRCm38.p2 assembly of mouse genome using STAR aligner^12^ with Gencode vM2^13^ transcripts included into pre-generated reference for reads 70 bp long (--sjdbOverhang 69). We then randomly downsampled the RNA-seq reads of each sample to generate datasets of 1M, 2M, 3M, 4M, 5M, 7.5M, 10M, 12.5M, 15M, 20M, 25M reads using fastq-sample utility from fastq-tools. Using these down-sampled sequence reads, we generated raw counts of number of tags on each gene by using htseq-count utility from HTseq package^14^. Gencode vM2 (with –t exon option) was used as a reference GTF, and the library was processed as unstranded (-s no). DESeq2 package^11^ (Version 1.4.5) was used to detect differentially expressed genes between two samples. The found DE genes for each evaluation set we defined as **E**. The 2200 found DE genes at 30M depth with the same adjusted p-value threshold of 0.05 we defined as the verification set **V** of truly DE genes. So the true positive genes we defined as TP = V ⋂ E, false positive genes FP = E \ (V ⋂ E), false negative genes FN = V \ (V ⋂ E), true negative TN = <all genes> \ (FP + FN + TP). Then we used the four benchmark criteria: sensitivity = TP/(TP+FN), precision = TP/(TP+FP), specificity = TN/(FP+TN), accuracy = (TP+TN)/(TP+FP+TN+FN).

### Fitting and normalization procedures

All the R functions and brief tutorial on their usage is provided in the supplemntary material. If a random variable X follows a double Pareto Lognormal (dPLN) distribution, then its logarithm follows the normal-Laplace (NL) distribution, which can be described as a convolution of Normal and Laplace distributions. Assuming that the RNA-Seq data we analysed from Bottomly et al. follow the dPLN distribution, we fitted natural logarithm of expression counts with NL probability density function^7^. Before fitting we applied a cutoff so that in a cumulative form of a distribution were kept values only above a chosen Y-axis threshold which was the same across all fitted distributions (Fig. 2c and Supplementary Fig. 2). To carry out NL normalization we used reference distribution of higher sequencing depth than the one we were normalizing. At first we found normalization coefficients individual for each gene in the distribution we were transforming. For each i^th^ gene, we found a corresponding x-value on the integral of the pdf function, fitted to its distribution – x_in_ on 5M (Fig. 2c). The point on the integral of pdf curve fitted to the reference distribution with the same y-value as x_i,n_ have x-value x_i,r_. So the normalization coefficient for the i^th^ gene is X_i_ = x_i,r_/x_i,n_. Then we multiplied logarithm of gene counts of the i^th^ gene by the normalization coefficient X_i_. To get the normalized RNA-Seq data table in count values we simply took exponent of the expression levels after normalization rounded to the nearest integer.

**Geometric mean averaging** was done as in DESeq, DEXSeq, DESeq2. The normalization parameters sij are considered constant within a sample and are estimated with the median-of-ratios method via formula

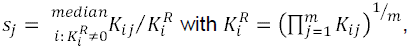

where K_ij_ is a read count for gene i in sample j^11^.

